# Why cancer cells have a more hyperpolarised mitochondrial membrane potential and emergent prospects for therapy

**DOI:** 10.1101/025197

**Authors:** Michael D Forrest

## Abstract

Cancer cells have a more hyperpolarised mitochondrial membrane potential (Ψ_IM_) than normal cells. Ψ_IM_ = ∼−220 mV in cancer cells as compared to ∼−140 mV in normal cells. Until now it has not been known why. This paper explains this disparity, in a mathematical framework, and identifies molecular targets and operations unique to cancer cells. These are thence prospective cancer drug targets. BMS-199264 is proposed as an anti-cancer drug. It inhibits the reverse, proton-pumping mode of ATP synthase, which this paper identifies as crucial to cancer cells but not to healthy, normal adult cells. In the cancer cell model, the adenine nucleotide exchanger (ANT) is inversely orientated in the mitochondrial inner membrane as compared to normal cells. This predicts it to have a different drug interaction profile, which can be leveraged for cancer therapy. Uncouplers, which dissipate the proton motive force, are proposed as anti-cancer medicines e.g. 2,4-dinitrophenol.

---

During aerobic respiration, the movement of electrons along the respiratory chain pumps protons across the inner mitochondrial membrane to build a proton motive force (pmf) [1-3]. The pmf is electrochemical, consisting of a hyperpolarised transmembrane voltage (Ψ_IM_, negative inside) and a proton concentration gradient (σpH, alkali inside). *In vivo*, with high concentrations of Pi, δpH is minor as compared to Ψ_IM_ [4]. Protons move down this electrochemical gradient, through ATP synthase, to generate ATP. Eukaryote cells *must* maintain a hyperpolarized voltage across their inner mitochondrial membranes. If this hyperpolarisation dissipates, the voltage-sensitive permeability transition pore (PTP) will open and release pro-apoptotic agents (e.g. cytochrome c) into the cytoplasm and drive apoptotic cell death [5].

In a normal cell, Ψ_IM_ flickers between -108 and -159 mV: with a mean value of -139 mV [6, 7]. Thermodynamically, the optimal Ψ_IM_ for maximal ATP production is between 130 to 140 mV, a rule that applies for all living organisms [7]. 10% value alterations in Ψ_IM_, above or below its optimum, results in a ∼90% decrease in ATP synthesis and a ∼90% increase in harmful reactive oxygen species (ROS) [7].

Cancer cells have a more hyperpolarised Ψ_IM_ than normal cells [8-17]. The more invasive and dangerous the cancer, the more hyperpolarised its Ψ_IM_ is observed to be [14-16]. The hyperpolarisation of Ψ_IM_ can be >50% greater in cancer cells than normal cells [14] e.g. Ψ_IM_ = ∼-210 mV in Neu4145 cancer cells [18]. The Ψ_IM_ hyperpolarisation in cancer cells can even be double that of normal cells [19]. So, generally, the Ψ_IM_ of cancer cells is extremely suboptimal for ATP production. However, these cancer cells aren’t using aerobic respiration and aren’t using Ψ_IM_ in the same way as normal cells.

Aerobic respiration is O_2_ dependent and uses glycolysis, the Krebs cycle and oxidative phosphorylation (OXPHOS) to produce ATP [1-3]. Aerobic glycolysis is the sole use of glycolysis to produce ATP, even in the presence of O_2_. Cancer cells can use aerobic glycolysis (Warburg effect) some or all of the time [18-39]. I propose that when in this mode, they have a more hyperpolarised Ψ_IM_. Indeed, experimentally, when cancer cells are switched out of aerobic glycolysis, into aerobic respiration, their Ψ_IM_ is returned to that of normal cells [18-19].

The fact that there is a disparity in Ψ_IM_ between normal and cancer cells is well established and has already been leveraged in human drug trials [40-41]. Delocalized lipophilic cations (DLCs) can cross membranes and their positive charge means they are drawn to, and accumulate in, the mitochondrial matrix (negative inside, because of hyperpolarised Ψ_IM_). Cancer cells have a more hyperpolarised Ψ_IM_ and so DLCs are more attracted to, and better retained by, their mitochondria than that of normal cells [12]. Using the Nernst equation [14], if the Ψ_IM_ of a cancer cell is 60 mV more hyperpolarised than that of a normal cell – which is within the range of observation [14, 18-19] – then a single charged DLC will accumulate 10 times more in the mitochondrial matrix of cancer cells than normal cells (T=300 K). DLCs with a double charge will accumulate 100 times more [42]. So, DLC poisons are more targeted to cancer cells and this means there are likely to be doses that can kill cancer cells, but leave normal cells unharmed. Different DLCs have been shown to accumulate in and selectively kill cancer cells, *in vitro* and *in vivo* [11, 43-47]. This affirms that cancer cells do have a more hyperpolarised Ψ_IM_, although no DLC has been successful in clinical trialling to date. For example, MKT-077 caused renal toxicity in Phase 1 trials [40-41].

It is not known why or how cancer cells have a more hyperpolarised Ψ_IM_. I provide a quantitative explanation, which identifies molecular targets and operations unique to cancer cells. These can be leveraged as cancer drug targets. To understand how Ψ_IM_ generation differs in cancer cells, we must first explain it for normal cells.

## The biophysics of Ψ_IM_ in normal cells [4, 48-50]

Mitochondrial ATP synthase (F_0_F_1_-ATPase) can synthesise or hydrolyse ATP. Protons can flow “downhill” through the ATPase, to generate ATP, or be pumped “uphill” by the ATPase, using ATP. The mitochondrial Adenine Nucleotide Transporter (ANT) can export ATP^4−^ for the import of ADP^3−^, or conduct the inverse. So, both ATPase and ANT catalyse reversible processes. Their directionality is governed by the mitochondrial membrane potential (Ψ_IM_) in relation to their reversal potential, E_rev_ATPase_ and E_rev_ANT_ respectively (mV). Which are set by the concentrations of the participating reactants, as shown in Equations 1-11 [4, 50].

## [1] The ATPase

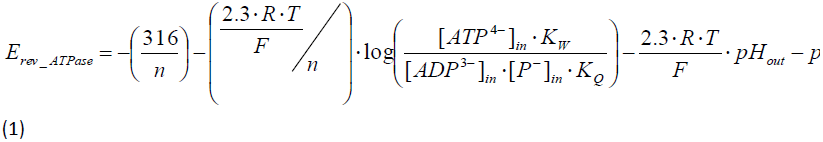

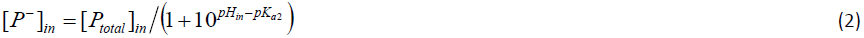

Where *in* relates to inside the mitochondrial matrix, *out* relates to outside the matrix (mitochondrial intermembrane space and cytoplasm), *n* is the H^+^/ATP coupling ratio of the ATPase, *R* is the universal gas constant (8.31 *J · mol*^−1^), *F* is the Faraday constant (9.64*10^4^ *C · mol*^−1^), *T* is temperature (K), K_W_ is the affinity constant for ADP, K_Q_ is the affinity constant for ATP, [*P*^−^]_*in*_ is the free phosphate concentration inside, [*P*_*total*_]_*in*_ is the total phosphate concentration inside, *pK*_*a*2_ = 7.2 for phosphoric acid (H_3_PO_4_).

## [2] The ANT exchanger

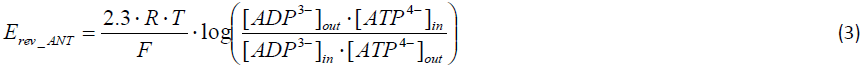

Note in Equations 1, 2 and 3: that the ATPase and ANT only (directly) share 2 common reactants: [*ATP*^4−^]_*in*_ and *ATP*^3−^]_*in*_.

## [3] The free ATP concentration inside [*ATP*^4−^]_*in*_

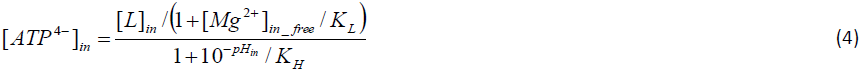

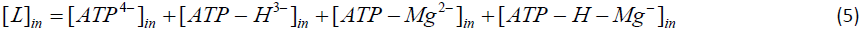

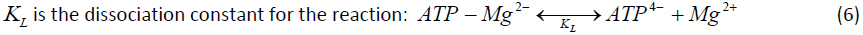

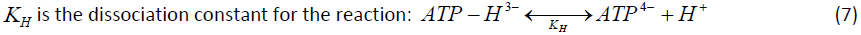

## [4] The free ADP concentration inside[*ADP*^3−^]_*in*_

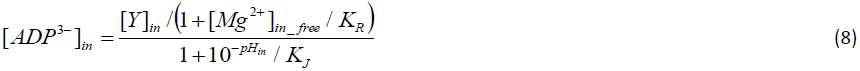

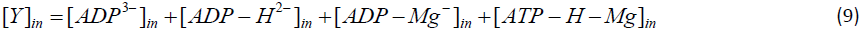

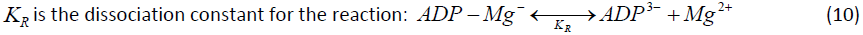

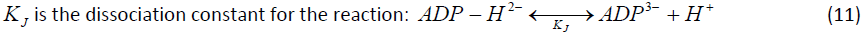

## [5] The free ADP and ATP concentrations outside – [*ADP*^3−^]_*out*_ and [*ATP*^4−^]_*out*_ respectively – are calculated by equations of the same form as those for inside; but are not shown here for brevity

The dissociation constants for [*ADP*^3−^]_*out*_ are K_Z_ and K_X_. The dissociation constants for [*ATP*^4−^]_*out*_ are KO and KU.

Figure 1 shows computational estimations of E_rev_ATPase_ and E_rev_ANT_ (mV) at different [ATP]_in_/[ADP]_in_ ratios [4, 48, 50]. This graph was made by computing Equations 1-11 with parameters from [4]: [ATP]_out_=1.2 mM, [ADP]_out_= 10 μΜ, [P]_in_= 0.01 M, n = 3.7 (2.7 for the ATP synthase plus 1 for the electrogenic ATP^4−^/ADP^3−^ exchange of the ANT and the nonelectrogenic symport of phosphate and a proton by the phosphate carrier [51]), pH_in_ = 7.38, pH_out_= 7.25, T = 310 K, [Mg^2+^]_in_free_=0.5 mM, K_W_= 10^−3.198^, K_Q_= 10^−4.06^, K_L_= 0.114 mM, K_H_= 10^−7.06^ M, K_R_= 0.906 mM, K_J_= 10^−6.8^ M, K_Z_= 0.906 mM, K_X_= 10^−6.8^ M, K_O_= 0.114 mM, K_U_= 10^−7.06^ M. Traces were computed by Erev estimator software [4, 48, 50], which can be downloaded at: http://www.tinyurl.com/Erev-estimator.

**Figure 1.**
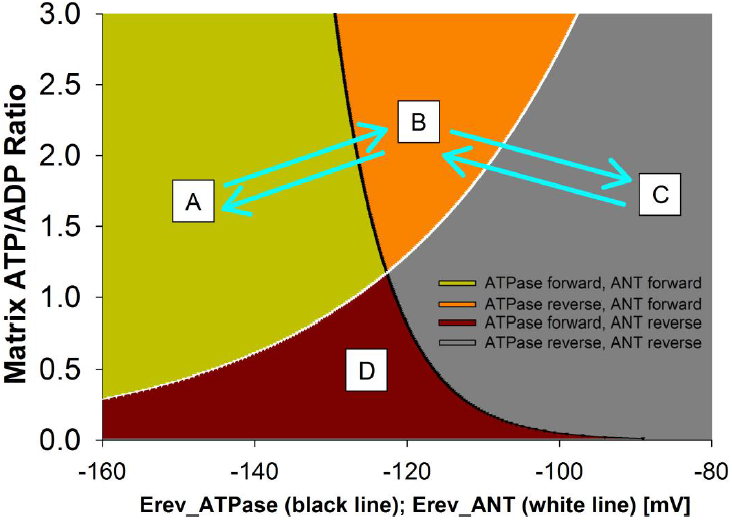
[4, 48, 50]; Computational estimations of E_rev_ATPase_ and E_rev_ANT_ (mV) at different mitochondrial matrix [ATP]_in_/[ADP]_in_ ratios. Black line is E_rev_ATPase_, white line is E_rev_ANT_. Note that a decreased [ATP]_in_/[ADP]_in_ ratio increases E_rev_ATPase_ and decreases E_rev_ANT_. A, B, C and D mark different operating states: (A) ATPase forward, ANT forward. (B) ATP reverse, ANT forward. (C) ATPase reverse, ANT reverse. (D) ATPase forward, ANT reverse.

During OXPHOS, protons are pumped by the complexes of the respiratory chain out of the mitochondrial matrix and into the mitochondrial intermembrane space. This hyperpolarises Ψ_IM_ and makes it more negative than E_rev_ATPase_ and E_rev_ANT_ (the green coloured “A-space” of Figure 1). With Ψ_IM_ hyperpolarised to E_rev_ATPase_, ATPase works in its “forward” mode and synthesises ATP. With Ψ_IM_ hyperpolarised to E_rev_ANT_, ANT works in its “forward” mode and exports mitochondrial matrix ATP for the import of cytoplasmic ADP. So, the mitochondrion produces and exports ATP. The “forward” operation of ANT and ATPase is a depolarising force to Ψ_IM_. ATP^4−^ export for ADP^3−^ import is depolarising and so are protons flowing “downhill” through ATPase. However, Ψ_IM_ doesn’t depolarise because at the same time protons are being continually pumped “uphill” by the respiratory chain complexes, which is a hyperpolarising drive to Ψ_IM_. Actually, Ψ_IM_ doesn’t remain constant during OXPHOS – but “flickers” (as much as >100 mV) [48-50] as these depolarising and hyperpolarising forces wrestle back and forth for a temporary *net* dominance.

If Ψ_IM_ is more positive (depolarised) than E_rev_ATPase_ and E_rev_ANT_, they both work in their “reverse” mode (the grey coloured “C-space” of Figure 1). ATPase hydrolyses ATP and ANT imports cytoplasmic ATP for the exchange of mitochondrial matrix ADP. So, the mitochondrion imports and consumes ATP. In this state, ATPase pumps protons into the intermembrane space which hyperpolarises Ψ_IM_. In addition, ANT imports ATP^4−^ and exports ADP^3−^, so a negative charge is gained on the matrix side which hyperpolarises Ψ_IM_.

During OXPHOS, Ψ_IM_ is more negative than E_rev_ATPase_ and E_rev_ANT_. If aerobic respiration is switched off, for example if the cell switches into aerobic glycolysis, then there will no longer be the hyperpolarising offset to the depolarising, “forward” action of ANT and ATPase: Ψ_IM_ will depolarise. E_rev_ATPase_ is more negative than E_rev_ANT_ and so Ψ_IM_ will depolarise past this reversal potential first. In this case, ATPase will switch into its “reverse” mode and ANT will remain in its “forward” mode (the orange coloured “B-space” of Figure 1). So, the ANT action will remain depolarising but the ATPase action will switch to being hyperpolarising – pumping protons out rather passing protons in. However, this reverse ATPase action requires ATP and with ANT pumping ATP out, there is little to be had. Furthermore, near the reversal potential of ATPase there is little driving force for an ATPase action. Hence, depolarising forces dominate and Ψ_IM_ depolarises further. When Ψ_IM_ is equal to E_rev_ANT_ then ANT does no “forward” or “reverse” ATP/ADP exchange and its effect on Ψ_IM_ is lost. At this point: With no ATP coming into the matrix, ATPase can no longer hydrolyse ATP to pump protons and its hyperpolarising action is also lost. In the absence of these forces, ensuing proton leak will depolarise the membrane potential. This will then make Ψ_IM_ more depolarised than E_rev_ANT_ and permit ANT to conduct a hyperpolarising exchange of ATP/ADP. The more depolarised it is past E_rev_ANT_, the more drive there is for this hyperpolarising exchange and the more that will occur. The resultant ATP entry will permit ATPase to conduct a hyperpolarising pumping of protons. So, there are hyperpolarising forces, conducted by ATPase and ANT, that come into play to prevent further depolarisation past E_rev_ANT_. They cannot hyperpolarise Ψ_IM_ to be more negative than E_rev_ANT_ because these forces are largely lost at this point, but they can prevent further depolarisation past this point. The result is that Ψ_IM_ will oscillate around E_rev_ANT_. So, at the loss aerobic respiration Ψ_IM_ will converge to E_rev_ANT_. Hence, at the loss of aerobic respiration there is a “safety net” of mechanisms to prevent the collapse of Ψ_IM_ and the mass consumption of cytoplasmic ATP through mitochondrial proton pumping by ATPase.

At certain mitochondrial matrix [ATP]in/[ADP]in ratios, it is possible for ATPase to be in “forward” operation and ANT to be in “reverse” operation (the whine coloured “D-space” of Figure 1). The former is depolarising, the latter hyperpolarising. However, it may be unlikely for mitochondria to have such a hyperpolarised Ψ_IM_ and a low matrix [ATP]_in_/[ADP]_in_ ratio. So, it may be that this part of the graph has no biological representation and can be discounted [4, 50]. The Ψ_IM_ in the “D-space” prompts ATPase to create ATP and ANT to import ATP. The ensuing rise in the [ATP]_in_/[ADP]_in_ ratio would push the system out of the “D-space” and into another area of the graph (Figure 1).

If proton pumping by the respiratory chain is stopped, but the Krebs cycle still persists, Ψ_IM_ will depolarise past E_rev_ATP_ but not all the way to E_rev_ANT_. The Krebs cycle can produce ATP (or GTP) in its succinyl CoA to succinate step. Once Ψ_IM_ is less negative than E_rev_ATP_, the ATP produced by the Krebs cycle may support the “reverse” ATP hydrolysing, proton pumping, hyperpolarising action of ATPase [4, 48-50]. This action will “hold” Ψ_IM_ in this range between E_rev_ATP_ and E_rev_ANT_. However, during aerobic glycolysis: both OXPHOS and the Krebs cycle are shunted. So, this situation will not apply in this case. As aforementioned, Ψ_IM_ should converge upon and oscillate around E_rev_ANT_.

IF-1 is a physiological protein, expressed by some tissues of some organisms, that inhibits the consumption of ATP by the F_0_F_1_-ATPase [4]. So, it prevents the “reversal” of the ATPase upon depolarisation of Ψ_IM_ past E_rev_ATP_. Its blockage isn’t complete, but increases with matrix [ATP], decreased matrix pH (acidification) and dissipated Ψ_IM_. With F_0_F_1_-ATPases unable to “reverse”, to confer a hyperpolarising pump of protons, they offer little resistance to an external, imposed depolarisation. So, this imposed depolarisation can converge relatively unopposed to E_rev_ANT_. Depolarisation past this point switches the ANT into producing a hyperpolarising exchange and this tries to “hold” Ψ_IM_ at E_rev_ANT_, as described earlier. The result is that Ψ_IM_ will oscillate around E_rev_ANT_. Unless the imposed depolarisation is strong enough to overcome this resistance, in which case the continued depolarisation will eventually open the voltage-dependent PTP and apoptosis is then all but assured.

## Why cancer cells have a more hyperpolarised Ψ_IM_ is mysterious

As aforementioned, I suggest that cancer cells have a more hyperpolarised Ψ_IM_ when they are utilising aerobic glycolysis. I suggest that this mode is a function of cancer proliferation and so the more aggressive and dangerous the cancer, the more time they spend in this operating state. During aerobic glycolysis, the Krebs cycle and OXPHOS are shunted and aren’t used. By the reasoning of the previous section, Ψ_IM_ should thus converge upon – and oscillate around – E_rev_ANT_. However, there is a problem. Refer to Figure 1 and note that E_rev_ANT_ is in a range around ∼−120 mV (−115 mV at matrix [ATP]_in_/[ADP]_in_ ratio = 1.5). But the Ψ_IM_ of cancer cells is much more hyperpolarised: e.g. the Ψ_IM_ of Neu4145 cancer cells is ∼-210 mV [18].

## Cancer cells use a different ANT isoform

The ANT referred to thus far in this manuscript is the ANT1 isoform. There are 4 human ANT isoforms (gene names in brackets): ANT1 (SLC25A4), ANT2 (SLC25A5), ANT3 (SLC25A6) and ANT4 (SLC25A31) [52]. They have ∼90% homology, except ANT4 which has ∼70% homology to the others. A comparative study in yeast with the heterologous expression of ANT1, ANT2 and ANT3 (underneath the same promotor region) showed them all to have similar adenine nucleotide exchange characteristics [52]. ANT1 is expressed in differentiated adult cells. ANT2 is expressed in rapidly proliferating cells (embryonic stem (ES) cells, cancer cells). ANT3 is expressed at a low level ubiquitously. ANT4 is highly expressed in male gametes.

In differentiated adult cells, ANT1 is expressed highly and ANT2 expression is minimal [53]. I surmise that as a differentiated cell turns cancerous, and as it switches from aerobic respiration to aerobic glycolysis, it downregulates ANT1 and upregulates ANT2 expression. Indeed, in the majority of cancer cell lines, ANT1 expression is very low, whereas the expression of ANT2 is very high [53]. ANT3 expression is low across the board.

ANT1 and ANT3 are associated with aerobic respiration. They export the ATP produced by OXPHOS from the mitochondria into the cytosol, while importing ADP [54]. ANT2, by contrast, is associated with aerobic glycolysis [54]. When a cell is rapidly proliferating (e.g. cancerous) it switches off OXPHOS and ANT2 may import glycolytically produced ATP into mitochondria, while exporting ADP [54]. “Reverse” ATP synthase action hydrolyses this ATP to pump protons and maintain Ψ_IM_ and so prevent apoptosis [54]. ANT2 expression is a marker for rapid proliferation and/or cancer. Hence, specific inhibition of ANT2 is a prospective anti-cancer strategy.

## But ANT2 may not import ATP into mitochondria

ANT2 import of ATP into mitochondria [54] is a hypothesis, although it is stated as an established fact in some of the literature, and there is a problem with this account [55-56]. Inhibition of OXPHOS alone (e.g. using myxothiazol, a Complex III respiratory inhibitor) cannot collapse the Ψ_IM_ of a cell using aerobic respiration. In addition, ATP synthase must be inhibited (e.g. using oligomycin) or ANT must be inhibited (e.g. using carboxyatractyloside or bongkrekic acid). Inhibiting ATP synthase or ANT inhibits the “reverse” action of ATP synthase, which can maintain Ψ_IM_ in the absence of OXPHOS. ANT inhibition services this requirement by cutting off ATP delivery to ATP synthase. However, this is all the case for cells using aerobic respiration. It is not the case for cancer cells. In A549 human lung cancer cells, myxothiazol slightly decreases Ψ_IM_ and subsequent oligomycin collapses Ψ_IM_. However, carboxyatractyloside and bongkrekic acid fail to collapse Ψ_IM_. Nonetheless, subsequent oligomycin does lead to full depolarization. Similarly, 2-deoxyglucose, a glycolytic inhibitor, can collapse Ψ_IM_. These results show that the cancer cells are using mitochondrial ATP synthase to hydrolyse glycolytic ATP, to maintain Ψ_IM_. But that in cancer cells, entry of glycolytic ATP into mitochondria can occur by a pathway other than ANT [55]. This pathway is unknown. What could it be?

It could be through the electroneutral ATP-Mg/Pi carrier (APC) [55]. APC exchanges ATP (Mg-ATP^2−^) for Pi (HPO4^2−^) [57-58]. APC can alternatively exchange ADP (HADP^2−^) for Pi, if Mg^2+^ is absent. So, it could perform some “fudged” electroneutral ATP-ADP exchange.

Some intracellular parasites express ATP transporters on their plasma membrane when inside a host cell, to steal ATP from the host’s cytoplasmic pool. There is a diversity of such parasites – e.g. chlamydia and rickettsiae bacteria [59-61], *Lawsonia intracellularis* [62] and the eukaryote: *Encephalitozoon cuniculi* [63]. PamNTT1, the ATP/ADP transporter from the amoeba symbiont *Protochlamydia amoebophila* (a chlamydia-related bacterium) [59] has an electroneutral action independent of the membrane potential. It exchanges [ATP^4−^] *in* for [ADP^3−^ and Pi^−^] *out* [63]. So, it is very distinct from ANT. Furthermore, it likely has 11-12 transmembrane domains; which is a significant difference from the 6 transmembrane helices of ANT and other members of the mitochondrial carrier family (MCF). I used BLAST [64] at http://blast.ncbi.nlm.nih.gov, with its default search parameters for “somewhat similar sequences” (blastn algorithm), to search the human genome for a PamNTT1 (NCBI accession number: AJ582021) homologue. With the idea that maybe such a homologue could be how rapidly proliferating/cancer cells import ATP into their mitochondria. However, none was found (data not shown). I then used BLAST again, with the same search parameters, to search the human genome for homologues of the ATP/ADP transporters from the eukaryote *E. cuniculi* [65]: EcNNT1–4 (GenBank accession numbers: EU040266-EU040270). However, none was found (data not shown).

Members of the mitochondrial carrier family (MCF) have a tripartite structure. That consists of three homologous sequence repeats of about 100 amino acid residues. Which each have a signature motif: P-X-[D/E]-X-X-[R/K] [66] (Prosite: PS50920). ANT and APC are members. The human genome likely encodes 48 different mitochondrial carriers [66]. A sizable proportion of these have not yet had their function assigned. It could be that one or more of these “orphan” transports facilitate ATP entry into mitochondria.

## ANT2 is essential to cancer cells; ANT1 and ANT3 kill cancer cells

It might still be that ANT2 transports glycolytic ATP into mitochondria in cancer cells, but it isn’t the only pathway for this. There may be a redundancy in this cancer system. Or ANT2 may have some other role. ANT2 does seem crucial to cancer cells.

I propose that cancer cell metabolism is similar to that of embryonic stem (ES) cells. Indeed, they share genetic expression fingerprints [67-68] and ES cells have a hyperpolarised Ψ_IM_ also [69]. They both employ aerobic glycolysis some or all of the time [18-39, 70], are immortal (divide forever without limit) [71-72] (as a function of using aerobic glycolysis [73]), respond to ROS damage by apoptosis rather than repair [19, 74] and can proliferate rapidly. So, with caution, we can learn more about cancer from ES cells and vice versa. In mice, ANT2 deficiency is embryonically lethal [75]. ANT2 is crucial to ES cells and we extrapolate from this to suggest that it is crucial to cancer cells. Indeed, ANT2 knockdown (RNA interference, shRNA) represses cancer proliferation and induces apoptotic death to cancer cells *in vitro* and *in vivo* [76]. Although others have reported ANT2 knockdown to have no such effect [77]; but this earlier, alternative report can be considered an inferior study because it used siRNA rather than shRNA; shRNA produces a more complete, robust, long lasting, long term knockdown [76].

In cancer cells, whereas ANT2 is anti-apoptotic [76], ANT1 is pro-apoptotic [78]. Overexpression of ANT1 induces apoptosis in cultured cancer cells by collapsing Ψ_IM_ and opening the voltage-dependent PTP [78]. Indeed, ANT1 transfection significantly suppresses tumor growth *in vivo* [78]. ANT3 is pro-apoptotic also [79]. Interestingly, over-expression of ANT1 is lethal to embryo cells [80], like it is to cancer cells.

## I predict that ANT2 is inserted in the inner mitochondrial membrane in the opposite orientation to ANT1 and ANT3

I suggest that ANT2 ***does*** import ATP into mitochondria in cancer cells (and ES cells). I predict that ANT2 is orientated in the inner mitochondrial membrane in the opposite orientation than ANT1 and ANT3 (Figure 2). Furthermore, that this makes ANT2 resistant to inhibition by carboxyatractyloside (CAT) and bongkrekic acid (BKA), which are both inhibitors of ANT. This accounts for why these drugs cannot block ATP entry into the mitochondria of cancer cells, as mentioned earlier and described in [55-56].

**Figure 2.**
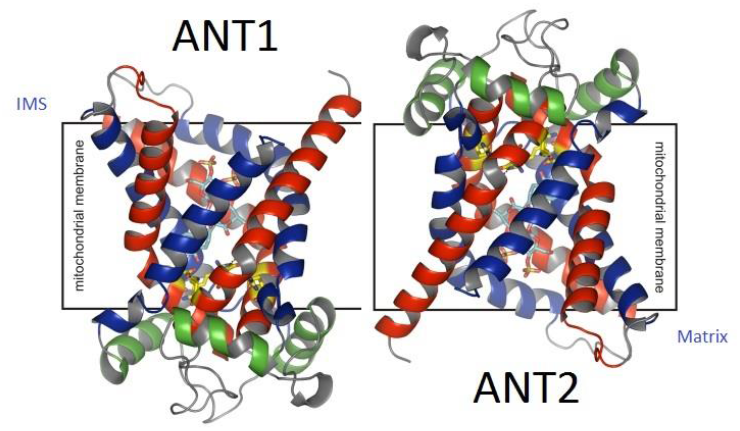
Orientation of ANT1 [84] and the predicted, inverse orientation of ANT2. The mitochondrial intermembrane space (IMS) and matrix are labelled.

BKA can be lipid soluble and can cross the inner mitochondrial membrane. CAT isn’t and can’t [81]. BKA crosses membranes as the electroneutral BKAH3. But it is its anionic form, BKA^3−^, which inhibits ANT [81]. CAT binds the intermembranous (“cytoplasmic”) side of ANT. BKA binds the matrix side of ANT [81]. If ANT2 is inserted in the inner mitochondrial membrane in the opposite orientation than ANT1/ANT3 – if its “m-side” is instead facing the cytoplasm (intermembranous space) and its “c-side” is instead facing the mitochondrial matrix – I anticipate that it will be resistant to inhibition by CAT or BKA. CAT won’t be able to cross the inner mitochondrial membrane to access its binding site on the “c-side” of ANT, which is now in the mitochondrial matrix. BKA will be able to access its binding site on the “m-side” of ANT. But now this site is located in the acidic, proton rich environment of the intermembrane space. So, here, BKA^3−^ will quickly pick up protons and be converted to BKAH_3_. This form cannot inhibit ANT.

If ANT2 is orientated oppositely to ANT1/ANT3, it could confer it very different adenine nucleotide exchange kinetics; which are more suitable for cancer cells. Indeed, it may favour the import, rather than the export, of ATP. Without this re-interpretation, it is hard to see why ANT2 is so vital to cancer cells [76], and ANT1 and 3 so harmful [78-79], while they all have high sequence homology (∼90%). And similar adenine nucleotide transport capabilities when expressed in a heterologous yeast system [52, 82]. I surmise that this heterologous system doesn’t have the infrastructure to orientate ANT2 oppositely. So, ANT1, ANT2 and ANT3 are all orientated in the same way and, in this scenario, they do all have similar exchange characteristics. A further distinction is that an opposite orientation of ANT2 may make it less predisposed to joining in to make the permeability transition pore (PTP) in apoptotic scenarios (ANT can be a PTP component [83]).

An opposite orientation of ANT2 opens prospects for selective drugs. A drug that binds the “m-side” of ANT molecules, but that can’t cross the inner mitochondrial membrane, may inhibit ANT2 (which has its “m-side” facing the cytoplasm) but not ANT 1 or 3 (which have their “m-side” in the mitochondrial matrix). So, this would be a selective anti-cancer drug. Alternatively, a drug (lipid or non-lipid soluble) that only binds the “m-side” of ANT in an acidic environment might selectively inhibit ANT2. It has its “m-side” in the acidic intermembranous space (IMS) rather than the alkali mitochondrial matrix. A delocalised lipophilic cation (DLC), that binds the “c-side” of ANT, would be preferentially located to the mitochondrial matrix by Ψ_IM_ where it could inhibit ANT2 and not ANT1/3. Alternatively, a drug that binds the “c-side” of ANT, tethered to a lipophilic cation to convey it passage through the inner mitochondrial membrane, and a positive charge(s) for targeting to the mitochondrial matrix, could inhibit ANT2 and not ANT1/3. A lipophilic anion (or a molecule tethered to one), that binds the “m-side” of ANT, would be preferentially located to the mitochondrial IMS by Ψ_IM_ where it could inhibit ANT2 and not ANT1/3. Antibodies against the “m-side” or “c-side” of ANT may yield novel inhibitors. That could then be used in cancer therapy, to leverage the orientation disparity between ANT2 in cancer cells and ANT1/3 in normal cells. We have an ANT crystal structure [84]. The “m-side” and “c-side” of ANT are now conceivable cancer drug targets and should be targeted via *in silico* virtual screening, structure based drug design and high-throughput *in vitro* screening e.g. using chip technologies [85].

## The E_rev_ANT_ of ANT1 and ANT3 is dangerous for cancer cells and this is why they downregulate them

ANT3 is pro-apoptotic to cancer cells [79], ANT1 also [78]. Over-expression of ANT1 induces apoptosis in cultured cancer cells by collapsing Ψ_IM_ and opening the voltage-dependent PTP [78]. For normal adult cells, we previously discussed how the E_rev_ANT_ of ANT1 protects Ψ_IM_ from depolarising to a voltage that opens the PTP and causes apoptosis. But here, in cancer cells, we see that ANT1 is pro-apoptotic [78] rather than anti-apoptotic as it is in normal cells [4, 48-50]. I suggest that this is because the E_rev_ANT_ of ANT1 is more depolarised in cancer cells. If Ψ_IM_ converges to, and is held at this new E_rev_ANT_ potential, far from being protective – this takes Ψ_IM_ to a depolarised potential that opens the PTP and brings apoptosis. It is for this reason that cancer cells heavily downregulate the expression of ANT1 (and ANT3); such that it has a minimal effect on cancer metabolism unless it is upregulated by an artificial/experimental/therapeutic intervention. Why is the E_rev_ANT_ of ANT1 (and ANT3) more depolarised in cancer cells?

A high cytosolic ATP/ADP ratio inhibits glycolysis, through allosteric feedback of ATP on key glycolytic enzymes [1]. In aerobic respiration, much of this ATP will come from mitochondrial export. During aerobic glycolysis, ATP is imported into, rather than mass-exported out of, mitochondria and the ATP/ADP ratio is much lower in the cytoplasm [55]. A low cytoplasmic ATP/ADP ratio favours a high glycolytic rate, which is needed for rapidly proliferating cells. Indeed, one of the postulated reasons as to why rapidly proliferating cells favour aerobic glycolysis (with its higher glycolytic rate) some or all of the time is because glycolytic intermediates are needed in large quantities for macromolecular biosynthesis [86-87]. A lower cytosolic ATP/ADP ratio shifts the E_rev_ANT_ of ANT1 (and ANT3) to more depolarised potentials. Figure 3(B) shows the shift in E_rev_ANT_ when cytoplasmic [ATP] is reduced fivefold from 1.2 to 0.24 mM and cytoplasmic [ADP] is increased fivefold from 10 to 50 μM (so the cytoplasmic ATP/ADP ratio decreases 25-fold). The midpoint on the E_rev_ANT_ curve, taken where the mitochondrial matrix ATP/ADP ratio on the y-axis=1.5, shifts from −115 to −30 mV. Incidentally, these changes do not alter the E_rev_ATP_ curve.

**Figure 3.**
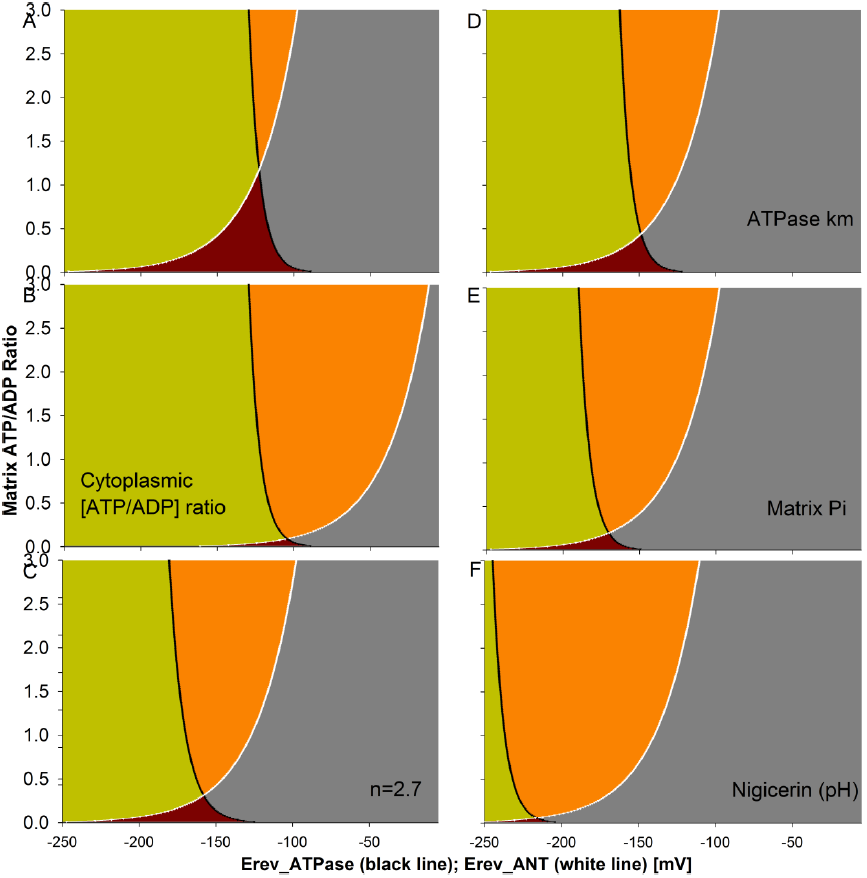
Computational estimations of E_rev_ATPase_ (black line) and E_rev_ANT_ (white line), which show their dependence on certain key parameters. (A) Same plots, using the same parameters, as Figure 1 (but shown with a different x-axis range). (B) Cytoplasmic [ATP/ADP] ratio changed from 120 to 4.8. (C) ATPase coupling ratio, *n*, changed from 3.7 to 2.7. (D) ATPase affinity for ADP and ATP modified: *K*_*w*_ changed from 10^−3.198^ to 10^−1.599^, *K*_*q*_ from 10^−4.06^ to 10^−6.09^. (E) Matrix Pi level reduced from 10 to 0.01 mM. (F) Matrix pH decreased from 7.38 to 6.38, intermembrane space pH increased from 7.25 to 8.25. This represents the action of the nigericin ionophore.

If there is a lower cytosolic ATP/ADP ratio we might surmise that there is a higher mitochondrial matrix ATP/ADP ratio. In Equations 1-11, a higher matrix ATP/ADP ratio shifts the E_rev_ANT_ of ANT1 (and ANT3) to more depolarised potentials. Looking at the y-axis of Figure 3(B), if the ATP/ADP matrix ratio=1.5: E_rev_ANT_ is −30 mV. If the matrix ratio=3, E_rev_ANT_ is −12 mV. Increasing the matrix ATP/ADP depolarises E_rev_ANT_ but it hyperpolarises E_rev_ATP_. At matrix ratio = 1.5, E_rev_ATP_ is −125 mV. At matrix ratio=3, E_rev_ATP_ is −130 mV.

## The E_rev_ATP_ of ATP synthase is more hyperpolarised in cancer cells

The coupling ratio of ATP synthase, *n*, is how many protons need to flow “downhill” for it to synthesise one ATP molecule from ADP and Pi. 2.7 protons need to flow through ATP synthase itself but the final value is one proton more than this: 3.7 [51]. This is because a proton is needed by the mitochondrial phosphate carrier to symport one Pi molecule into the mitochondrial matrix. Equation 1 employs this *n* parameter and *n*=3.7 in Figure 1. The coupling ratio is widely believed to be equal for the “forward” and “reverse” modes of ATP synthase. That is, in the “reverse” mode, the hydrolysis of one ATP molecule can pump the same number of protons that are required to synthesise one ATP molecule in the “forward” mode. I suggest that this isn’t true. The 2.7 value ascribed to ATP synthase itself likely holds. However, in the “reverse” mode there is no longer the need for Pi import and its associated proton cost. So, this renders *n*=2.7. Actually, Pi needs to be exported and this can be done by the ATP-Mg/Pi carrier (APC) and/or the phosphate carrier, which has an alternative mode that can perform an electroneutral exchange of a Pi molecule for a hydroxyl ion (OH^-^). The latter could be suspiciously stretched to be considered proton import if one considers a hydroxyl ion, with its proton component, equivalent to a proton flow. However, the former definitely cannot be. Figure 3(C) shows the more hyperpolarised E_rev_ATP_ plot when *n*=2.7 instead of 3.7. The midpoint of the E_rev_ATP_ curve (where matrix ATP/ADP ratio = 1.5) shifts from −125 to −174 mV. If *n*=2.2, the midpoint shifts to −215 mV (not shown). Incidentally, the E_rev_ANT_ plot is unchanged.

ANT2 expression is under the control of the glycolysis regulated box (GRBOX), which regulates expression of machinery for the aerobic glycolysis operating state [53-54]. Also under this control is a gene for a β subunit of ATP synthase [53]. The F_1_F_0_ ATP synthase consists of two domains: a transmembrane proton translocating F_0_ domain (with subunits ab_2_c_12_) and a cytoplasmic ATP catalytic domain (with subunits α3β3γδε) [88]. The two domains are attached by a “stalk” of subunits γ and ε. This new β subunit in the catalytic domain of ATP synthase may convey a different affinity constant, *Km*, for ADP and/or ATP (*K_w_* and *K*_*q*_ in Equation 1 respectively). Increasing *K*_*w*_ and/or decreasing *K*_*q*_ hyperpolarises E_rev_ATP_. Increasing *K*_*w*_ by 50% (from 10^−3.198^ to 10^−1.599^) and decreasing *K*_*q*_ by 50% (from 10^4.06^ to 10^−6.09^) shifts the midpoint of the E_rev_ATP_ curve from −125 to −185 mV, Figure 3(D). As aforementioned, mitochondria in cancer cells export Pi and no longer have its import, driven by the pmf. So, the Pi level in the mitochondrial matrix is likely to be lower than in normal cells. If [P]_in_ is reduced hundred-fold from 10 to 0.1 mM, as in Figure 3(E), then the midpoint of the E_rev_ATP_ curve shifts from −125 to −158 mV. The E_rev_ANT_ plot is unchanged.

## E_rev_ANT_ (ANT1/ANT3) and E_rev_ATP_ in cancer cells

Figure 4 shows these values modelled for cancer (panel B) and normal cells (panel A) using Equations 1-11. As compared to the normal cell parameters, the cancer equations have a fivefold reduction in the cytosolic ATP/ADP ratio ([ATP]_out_ is maintained at 1.2 mM, [ADP]_out_ is increased fivefold from 10 to 50 μM), the H^+^/ATP coupling ratio: *n* = 2.7 instead of 3.7, matrix Pi is reduced twentyfold from 10 to 0.5 mM and the matrix ATP/ADP ratio is doubled (*if* one reads the value on the x-axis at y-axis values: y = 3 for cancer panel and y =1.5 for normal panel). These cancer values are set by the rationale presented thus far and were chosen to present how the same equations used for normal cells can replicate the Ψ_IM_ of cancer cells, all be it with different parameter values. In Figure 4, E_rev_ANT_ = −115 mV, E_rev_ATP_ = −124 mV for normal (A); E_rev_ANT_ = −55 mV, E_rev_ATP_ = −211 mV for cancer (B). So, the cancer system has a more depolarised E_rev_ANT_ (for ANT1/ANT3) and a more hyperpolarised E_rev_ATP_. ANT2 is discussed in the next section.

**Figure 4.**
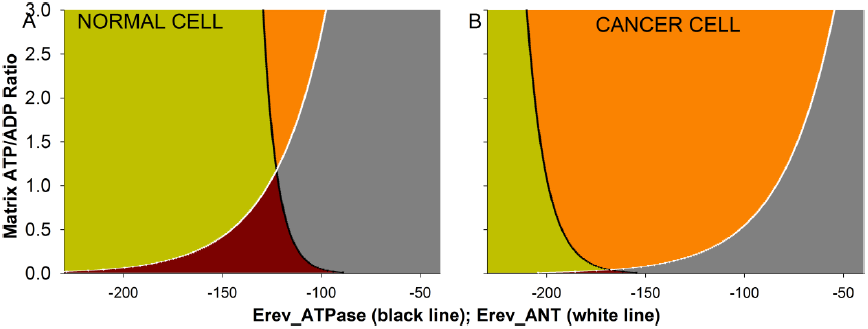
Computational estimations of E_rev_ATPase_ (black line) and E_rev_ANT_ (white line) for a normal cell (A) and a cancer cell (B). (A) Normal cell. Same plots, using the same parameters, as Figure 1 (but shown with a different x-axis range). (B) Cancer cell. Cytoplasmic [ATP/ADP] ratio decreased from 120 to 24, H^+^/ATP coupling ratio: *n* = 2.7 instead of 3.7, matrix Pi reduced from 10 to 0.5 mM and the matrix ATP/ADP ratio is doubled (*if* one reads the value on the x-axis, to get the E_rev_ value, at y-axis values: y = 3 for cancer panel and y = 1.5 for normal panel). E_rev_ANT_ = −115 mV, E_rev_ATP_ = −124 mV for normal cell (A); E_rev_ANT_ = −55 mV, E_rev_ATP_ = −211 mV for cancer cell (B).

## What is the E_rev_ANT_ of ANT2?

Firstly, note that ADP/ATP exchange by any ANT isoform is not energy dependent. It proceeds with high activity when the mitochondria are completely depolarised by uncouplers; in which case ADP and ATP are transported in both directions at nearly equal rates [81]. With the orientation of ANT1 and ANT3, a hyperpolarised Ψ_IM_ (negative inside) favours the export of ATP^4−^ and import of ADP^3−^. How energisation of the membrane affects the exchange by ANT2, in its postulated inverse orientation, is not known. The exchange of ANT1 in an inverse orientation has been studied in sub-mitochondrial vesicles produced by sonication. The vesicles typically form “inside-out” with their mitochondrial matrix face exposed on the outside of the vesicle [89]. In this inverted orientation, the exchange seems to occur without influence by the membrane potential (isn’t altered by uncoupling chemicals). But then there is the complication that how much membrane potential do these vesicles actually retain or sustain?

I think it is fair to suggest, in the absence of direct data, that at membrane potentials that ANT1 and ANT3 – in their “conventional” orientation – are exporting ATP: ANT2, in its “inverted” orientation, is importing ATP.

## The biophysics of Ψ_IM_ in cancer cells

In cancer cells Ψ_IM_ is hyperpolarised at ∼-220 mV. I suggest because E_rev_ATP_ is ∼-220 mV in cancer cells. And that Ψ_IM_ oscillates around this point. When more depolarised, ATPase is in it “reverse” mode and pumping protons at the expense of ATP hydrolysis to ADP and Pi. ANT2 imports ATP to service this hydrolysis and exports the ensuing ADP. The ATP-Mg/Pi carrier (APC) imports further ATP and exports Pi. Glycolysis in the cytoplasm synthesises ATP from this ADP and Pi. APC is electroneutral. The “reverse” mode of ATPase is hyperpolarising as is the ATP^4−^ import and ADP^3−^ export by ANT2. These hyperpolarising forces drive Ψ_IM_ to E_rev_ATP_ and then past it at which point ATPase switches into its depolarising “forward” mode. This then depolarises Ψ_IM_ back towards E_rev_ATP_. So Ψ_IM_ oscillates around E_rev_ATP_ (∼-220 mV). At E_rev_ATP_ precisely there is no drive for proton conductance or pumping through ATPase, so oscillating around this point ensures little ATP generation but not much ATP hydrolysis either. It is a “cheap” way to hold Ψ_IM_ at a hyperpolarised potential, safely well away from “dangerous” depolarised potentials that could open PTP and drive apoptosis. In cancer cells, ANT1 and ANT3 are expressed at low levels and so are irrelevant. However, if by an experimental intervention they are expressed at significant levels, they can kill the cancer cell. In cancer cells, the E_rev_ANT_ of ANT1 and ANT3 are abnormally depolarised. This means that their depolarising “forward” mode of operation – ATP^4−^ export, ADP^3−^ import – depolarises Ψ_IM_ towards their “dangerously” depolarised E_rev_ANT_ value. What is more, their export of ATP undermines the import of ATP by ANT2 and denies it to ATPase. Hence the hyperpolarising “reverse” mode of ATPase, wherein it needs ATP to pump protons, is compromised. Thus, it isn’t able to combat the depolarisation conveyed by ANT1 and/or ANT3. They have such depolarised E_rev_ANT_ values that the driving force for their depolarising exchange is immense at even rather modestly hyperpolarised potentials e.g. -100 mV.

## An alternative biophysics of Ψ_IM_ in cancer cells

The prior account assumes that cancer mitochondria are looking to maintain Ψ_IM_ by the lowest, “cheapest” ATP spend. However, an alternative view is that during aerobic glycolysis mitochondria are tasked with burning through ATP as a specific aim in itself [90]. To lower [ATP] in the cytoplasm, release key glycolytic enzymes from negative feedback by ATP [1, 90]), and permit high glycolytic rates. In proliferating cells, glycolysis isn’t just tasked with ATP production. It has important roles in shuttling its metabolic intermediates into macromolecular biosynthesis pathways [86-87]. The rapidly proliferating cell may have to forsake ATP to facilitate this [90]. Low cytosolic and high mitochondrial ATP/ADP ratios produce a hyperpolarised E_rev_ATP_ and depolarised E_rev_ANT_ for ANT1 and ANT3 (which are expressed at low levels). Proton pumping by ATP synthase (hyperpolarising), fuelled by ATP entry through ANT2 (hyperpolarising) and the ATP-Mg/Pi carrier (electroneutral), may be balanced electrically by ATP export/ADP import (depolarising) and/or proton leak (depolarising) by ANT1 and/or ANT3. ANT1 seems to convey a basal proton leak that ANT2 doesn’t [91-93]. This electrical balance (at ∼-220 mV) may keep Ψ_IM_ consistently and sufficiently depolarised to E_rev_ATP_ (E_rev_ATP_ < -220mV) such that ATP hydrolysis occurs at high rates. The amount of ANT1 and ANT3 is low but at this point of balance, Ψ_IM_ is sufficiently hyperpolarised to their E_rev_ANT_ that they produce crucial activity, despite being expressed at low levels. This is all a balance and will naturally have its fluctuations but if ANT1 and/or ANT3 are overexpressed: it depolarises Ψ_IM_ out of this balance and into a spiral of depolarisation, PTP opening and apoptosis. But then similarly if ANT1 (particularly) is underexpressed it will disturb the balance and Ψ_IM_ will hyperpolarise and oscillate around E_rev_ATP_ (the first scenario) and there won’t be as much ATP hydrolysis. So, this second account details a role for ANT1 and/or ANT3 in aerobic glycolysis where the first one rendered them redundant. It suggests that not just overexpression of ANT1 will perturb function, but under-expression of ANT1 will disturb function also. Indeed, ANT1 overexpression is pro-apoptotic [78] and ANT1 under-expression can kill cancer cells also [94]. In the latter case, cancer cells appear to die because of oxidative damage [94]. I postulate that ANT1 loss renders a lower rate of ATP hydrolysis in mitochondria and thence the glycolytic rate in the cytoplasm is lower, because of ATP feedback. A postulated role of enhanced glycolysis is protection from oxidative damage [73]. Indeed it conveys so much protection that it can gift cancer cells immortality [73]. A lower glycolytic rate conveys less protection and death. A high glycolytic flux permits a high flux into the pentose phosphate pathway (PPP) that branches from glycolysis. It produces NADPH from NADP+, which is needed for glutathione (GSH)-dependent anti-oxidant mechanisms. To protect, GSH needs to be in its reduced form and NADPH puts it into this reduced form (as it is converted to NADP^+^). GSH is needed by glutathione peroxidase (GP), which converts hydrogen peroxide (a ROS) into water. Upstream, superoxide dismutase (SOD) converts superoxide (O_2_•–, a ROS) into hydrogen peroxide. Increased GP activity will pull through greater SOD activity. So, less ANT1 produces less NADPH, less oxidative protection and death (by paraptosis). This is a different interpretation than that in the paper itself [94]. So, an imposed hyperpolarisation (e.g. by ANT1 knockdown) of the Ψ_IM_ in cancer cells may be able to kill them, in addition to an imposed depolarisation (e.g. by ANT1 overexpression).

## Re-interpreting the data of [94]

We both agree death is by oxidative damage. [94] suggests that lower ANT1 generates more ROS. I suggest, instead, that it compromises the enhanced ROS mitigation apparatus of cancer cells, by reducing their spurious ATP hydrolysis, thence their glycolytic rate, PPP rate and NADPH production. [94] talk of electron fumbles by the respiratory chain generating ROS, but during aerobic glycolysis electrons don’t passage along this chain. The higher ANT2 level in these cancer cells suggests they are utilising aerobic glycolysis; as does the lack of response to atractyloside (ATR) or bongkrekic acid (BKA). These drugs would kill a cell using aerobic respiration. They inhibit ATP/ADP exchange by ANT1 (but not ANT2 because of its inverted orientation; a prediction of this manuscript). They observe siRNA knockdown of ANT2 to have no effect but then as aforementioned, a working knockdown of ANT2 can require shRNA [76].

They observe enhanced glucose consumption, and presumably glycolysis, 72 hours after ANT1 has been reduced (by siRNA interference). But this could be an after-effect once the increase in un-sequestered ROS has committed the cell to death. It could be due to the active processes of paraptosis (a form of programmed cell death; an active process) consuming ATP, lowering ATP levels and reducing its allosteric feedback on key glycolytic enzymes and thus permitting a higher glycolytic rate.

The basal proton leak function of ANT1 can work even when its nucleotide exchange function is blocked by carboxyatractylate (CAT) [93] (and so maybe also ATR or BKA). It is likely to be this proton-leak function, and its loss, that is important in this case. So, uncoupling may be a physiological feature not just of aerobic respiration (“uncoupling to survive theory” [95]), but aerobic glycolysis also.

However, in contrast to [94], ANT1 knockdown by siRNA did not cause cancer cell death in a different study by a different group in a different cancer cell line [77]. Furthermore, ANT1 genetic knockout animals can live to adulthood [92]. So, this suggests that it isn’t vital to aerobic glycolysis (in the embryonic stage) or aerobic respiration in adult cells. However, there may be developmental adaptations in these knockout animals that make them non-typical e.g. an up-regulation of ANT3. Also, the knockout may not be complete.

It is unclear whether ANT1 associated proton leak is by a specific pathway within the carrier. Or, since the proton conductance is not dependent on the function or turnover of ANT, it may occur at the protein-phospholipid interface. It is still unclear to what extent this particular “function” is tractable to pharmacology. It is interesting that ANT1 conveys a proton leak that ANT2 doesn’t [91-92], which could be related to the postulated inverse orientation of ANT2.

## Cancer cells reduce ROS at source and sink

I propose that during aerobic glycolysis, ROS are confronted at source and sink. OXPHOS is shunted leading to less ROS generation and the mitigation system is upregulated leading to more ROS mitigation. ROS generation is determined by the redox state of NAD+, while the NADP^+^ redox state is pivotal to antioxidant defence. As compared to normal cells, cancer cells decrease NADH and increase NADPH levels. The latter may carry over to confer greater protection if the cancer cell periodically switches into aerobic respiration. Investigators have reported cancer cells to have higher NADH levels than normal cells [97]. But their spectroscopy can’t discriminate between NADH and NADPH and I suggest they are actually observing higher NADPH levels in cancer cells. Indeed, later studies with a spectroscopy that can distinguish between these two species reports higher NADPH, rather than NADH, in cancer cells [98]. Cancer may be combatted by increasing NADH [34] and/or lowering NADPH. This could be achieved by transfecting cancer cells with a mutant lactate dehydrogenase (LDH) that uses NADPH rather than NADH. Such a form has been engineered for a prokaryote LDH [99]. In a prior paper, I propose exogenous NADH as a cancer medicine [34].

## UCP2 depolarises Ψ_IM_ past E_rev_ATP_ and primes greater ATP hydrolysis by ATP synthase

Uncoupling proteins (UCPs) passage protons through the inner mitochondrial membrane, dissipating their electrochemical potential as heat [3, 100]. UCP1 is implicated in thermogenesis. The role of its homologues – UCP2, UCP3, UCP4 and UCP5 – is less clear. If Ψ_IM_ becomes too great during OXPHOS, there is greater ROS production [95]. A postulated role of UCP2 is to reduce Ψ_IM_ in this scenario and decrease ROS production [101]. UCP2 can be activated by reactive oxygen species (ROS), perhaps forming part of a negative feedback mechanism that shunts excessive ROS production [96]. ROS can produce mutations, which can lead to cancer [100]. A lower UCP2 activity can confer a greater risk of cancer [101]. However, once cancer develops – and it is utilising aerobic glycolysis – there is evidence that it upregulates UCP2 activity above that of normal cells [101]. Indeed, it is overexpressed in many cancers [101-102]. I reason that UCP2 conveys depolarisation that keeps Ψ_IM_ depolarised to E_rev_ATP_, such that significant ATP hydrolysis can occur, the cytoplasmic ATP/ADP ratio is kept low, glycolytic rate is high, PPP rate is high, NADPH levels are high and ROS are significantly sequestered. Indeed, UCP2 silenced cancer cells show a more hyperpolarised Ψ_IM_, a greater cytoplasmic ATP/ADP ratio and less lactate release than control cancer cells [103]. I suggest a more hyperpolarised Ψ_IM_ (closer to E_rev_ATP_) means that ATP synthase cannot hydrolyse as much ATP: this is why the cytoplasmic ATP level is higher, the glycolytic rate is lower and less lactate is released. I would argue that these cells have a lower ROS mitigation capability. Indeed, suppression of UCP2 (siRNA) results in greater ROS levels and the induction of apoptosis in cancer cells (during hypoxia, so any OXPHOS effects are redundant) [104]. Furthermore, overexpression of UCP2 is anti-apoptotic for cancer cells (during hypoxia, so any OXPHOS effects are redundant) [104]. Many anti-cancer drugs exert their effect by generating ROS. UCP2 inhibition potentiates their effect [105], perhaps by undermining the cancer cell’s ROS mitigation apparatus. For example, genipin is a UCP2 inhibitor. It sensitizes cancer cells to cytotoxic agents by decreased proton leak and increased ROS levels [106]. Genipin has been reported to induce apoptotic cell death in cancer cells via increased ROS [107-109]. UCP2 expression is higher in a modified cancer cell line lacking mitochondrial DNA, which cannot use OXPHOS and respires fully by aerobic glycolysis; UCP2 inhibition by genipin impairs the tumorigenicity of this cell line [102]. When UCP2 is overexpressed in the parent cancer cell line, that still has mitochondrial DNA, it decreases its Ψ_IM_ and increases its tumorigenicity [102]. UCP2, in addition to conferring a proton leak, is involved in the switch into aerobic glycolysis [103, 109-110]. siRNA-mediated UCP2 knockdown leads to reversal of the glycolytic phenotype in some cancer cells [111]. So, we see a link between aerobic glycolysis and resistance to chemotherapy. Cisplatin, one of the most important chemotherapeutics found thus far, acts by inhibiting UCP2 (amongst other targets) [112].

UCP2 exports pyruvate, oxaloacetate and related C4 compounds from mitochondria, denying them to aerobic respiration and helping the switch to aerobic glycolysis [102, 103, 109]. Indeed, in quiescent human pluripotent stem cells, high levels of UCP2 expression prevent mitochondrial glucose oxidation, favouring aerobic glycolysis, whereas during cell differentiation, UCP2 is repressed and glucose metabolism is shifted toward mitochondrial oxidation [110]. The former can be applied to cancer cells with our construct of equivalence-in the use of aerobic glycolysis – between cancer and ES cells. Furthermore, ROS are substantially increased in UCP2 shRNA knockdown pluripotent stem cells and result in elevated apoptosis [110].

This UCP2 role in cancer cells gives two contrasting points of attack. One can reduce UCP2 activity to hyperpolarise Ψ_IM_, reduce ATP hydrolysis, increase unmitigated ROS levels and bring cancer cell death. An alternative tact is to increase UCP2 expression/activity – to increase its depolarising proton flow – to collapse the mitochondrial Ψ_IM_ in cancer cells and bring apoptosis. Low doses of the uncoupler, FCCP, can replicate the lower ROS and anti-apoptotic effect of UCP2 overexpression in cancer cells; higher doses of FCCP produce cell death [105]. This FCCP experiment yields insight into the two disparate paths to killing a cancer cell via modulation of UCP2 proton leak.

## A drug that hyperpolarises Ψ_IM_ will convey a specific anti-cancer action e.g. nigericin

The antibiotic nigericin is an ionophore that performs K^+^/H^+^ exchange [3]. In normal cells, nigericin decreases the δpH component of the pmf, which prompts a compensatory increase in the hyperpolarisation of Ψ_IM_ (∼30 mV) to maintain the pmf [113]. If it hyperpolarises Ψ_IM_ in cancer cells, and it does in HeLa cells [114], this may be a basis to its observed anti-cancer action *in vitro* and *in vivo* [115]. Ionomycin can also cause Ψ_IM_ hyperpolarisation (∼10 mV) [113] and this may be a basis to its observed anti-cancer action *in vitro* and *in vivo* [116]. I suggest that an ionophore species that hyperpolarises Ψ_IM_ will convey an anti-cancer action. Figure 3(F) shows how nigericin might cause a hyperpolarisation in Ψ_IM_ in cancer cells. Nigericin acidifies the matrix and correspondingly increases the pH of the intermembrane space. Figure 3(F) shows matrix pH = 6.38, instead of 7.38, and intermembrane space pH = 8.25, instead of 7.25. Given the pH parameters of Equation 1, this moves E_rev_ATP_ to a more hyperpolarised potential. This increases the drive for, and action of, the hyperpolarising, proton-pumping “reverse” action of ATPase. At the same time, depolarising forces are diminished because E_rev_ANT_ also moves to a more hyperpolarised potential. Hence Ψ_IM_ hyperpolarises; it moves closer to E_rev_ATP_ than previously, so reducing the amount of ATP hydrolysis, and it may even converge upon the new hyperpolarised E_rev_ATP_ value, which would dramatically reduce the amount of ATP hydrolysis. This reduces the glycolytic rate which decreases the NADPH mediated ROS mitigation apparatus of cancer cells which promotes ROS mediated apoptosis. Furthermore, it sensitizes the cancer to other drugs or means (e.g. radiation) that increase ROS levels.

## Uncoupling Cancer: a drug that can depolarise Ψ_IM_ will convey a specific anti-cancer action

Exogenous uncouplers transport protons across the inner mitochondrial membrane and dissipate the pmf as heat [3]. Eukaryotes must maintain a hyperpolarised Ψ_IM_ or they will undergo apoptosis [5]. Aerobic respiration hyperpolarises Ψ_IM_ as it produces ATP; Aerobic glycolysis consumes ATP to produce a hyperpolarised Ψ_IM_. Under the challenge of an uncoupler drug, the former is more sustainable than the latter. Cancer cells use aerobic glycolysis, some or all of the time, and so will be exquisitely sensitive to uncoupling drugs. Furthermore, we have shown that cancer cells may have a very delicately balanced Ψ_IM_ value, which if mildly depolarised can tip into a runaway depolarisation because of the excessively depolarised E_rev_ANT_ values (for ANT1 and ANT3) in cancer cells. By contrast, in normal cells, the E_rev_ANT_ values are at more hyperpolarised values that are protective and act against excessive depolarisation.

Of course, a cancer cell could switch out of aerobic glycolysis and start using aerobic respiration to counter the uncoupling threat. But no, they can’t. Not long term. There is growing evidence that cancer cells *must* use aerobic glycolysis for at least a component of their proliferation cycle: constitutively activating OXPHOS kills cancer cells [19, 34, 37,] or halts their proliferation [18, 38-39] via ROS production [19, 34].

Additionally, because cancer cells have a more hyperpolarised Ψ_IM_ than normal cells, uncouplers will be selectively targeted to, and accumulated by, cancer cells. Uncouplers are lipophilic and can be cations or anions [117]. They will be more targeted to the mitochondrial matrix and intermembrane space of cancer cells respectively. At these locations they are primed to uncouple. If we consider, for example, that Ψ_IM_ = ∼-220 mV in cancer and Ψ_IM_ = ∼-140 mV in normal cells. This is an 80 mV difference which, using the Nernst equation (T = 300 k), suggests that lipophilic uncouplers will be accumulated and retained by cancer cells ∼20 times more if they are single charged and ∼500 times more if double charged. These differentials are significant. They mean that cancer cells will be selectively targeted and that, crucially, they will sequester the poison from normal cells. The more aggressive the cancer, the more hyperpolarised its Ψ_IM_ [14-16] and the more it will be targeted.

A major problem in cancer therapy is that cancer cells can accrue DNA mutations which confer drug resistance. This is a problem for drugs that target or rely (e.g. for transport) on DNA encoded proteins. The cancer cell develops a DNA mutation that confers a change in the protein structure which means the drug can no longer interact with it in the same way. Protonophores do not interact with or rely on proteins to collapse Ψ_IM_ and kill cancer cells.

The cytoplasm is, and needs to be, neutral in normal and cancer cells [118-120]. Tumours are acidic, normal tissue is neutral [120-122]. This is likely because cancer cells, unlike normal cells, are using aerobic glycolysis and excreting lactate and protons through the monocarboxylate symporter (a promising cancer drug target). The more aggressive the cancer is, the more acidic its tumour [123]. So, cancer cells, unlike normal cells, must maintain their intracellular pH above their extracellular acidity and protonophores will shuttle protons, undermine this homeostasis and kill cancer cells; a prediction.

## Uncouplers can kill cancer cells

2,4-dinitrophenol (DNP) and FCCP are uncouplers. In cancer cells, they cause cell cycle arrest at low doses and apoptosis at higher doses, via depolarising Ψ_IM_ which opens PTP [124-125]. The uncouplers: moronone [126], CCCP [127], clusianone [128] and hyperforin [129] also kill cancer cells. So, uncouplers can kill cancer cells. However, the crux issue is: can they kill cancer cells whilst leaving normal cells unharmed?

*In vitro*, F16 – a lipophilic cation uncoupler (a DLC) – kills cancer cells but *not* normal cells [11]. *In vitro*, nemorosone – a lipophilic anion uncoupler – kills HepG2 cancer cells (∼75% cell death) but not non-cancer human embryonic kidney HEK293T cells to the same degree (∼10% cell death) [130]. So, there is a possible selectivity of action; which might be enhanced if cancer and normal cells were to be tested side-by-side in the same *in vitro* assay. Because cancer cells, with their greater affinity for a charged lipophilic protonophore (as previously discussed), may accumulate and sequester it from the normal cells. A DLC derivative of gallic acid, TPP^+^C_10_, can uncouple and selectively kill cancer cells *in vitro* and in a singenic mouse model [131].

Valinomycin depolarises Ψ_IM_; nigericin hyperpolarises Ψ_IM_ [3]. As aforementioned, depolarisation or hyperpolarisation of Ψ_IM_ may kill cancer cells and these drugs both demonstrate an anti-cancer activity [115, 132]. Together, valinomycin and nigericin can uncouple H^+^, while K^+^ cycles around the membrane [3]. This combined uncoupling activity should be tested against cancer cells. Coumarins comprise a structurally diverse group of natural compounds found in a variety of plant sources [133-135]. Some coumarin molecules (mammea A/BA, mammea A/BB) can reduce tumour weight by 83% in test animals by halting the cell cycle and inducing apoptosis selectively in cancer cells, by an unknown mechanism [133]. Possibly by uncoupling Ψ_IM_: mammea A/BB collapses the Ψ_IM_ of the *Leishmania amazonensis* parasite [134]. A different coumarin molecule (mammea E/BB), with an anti-cancer action, has been shown to be an anionic protonophore with an uncoupling potency equivalent to that of FCCP [135].

Uncoupling chemicals can shuttle protons alone (e.g. DNP) or in interaction with a transmembrane protein in the inner mitochondrial membrane e.g. ANT [136]. There are other, further conceivable mechanisms to uncouple e.g. neutralising a negative molecular species residing in the mitochondrial matrix, shuttling a negative species out of the matrix, stimulating UCP activity or the intrinsic, basal uncoupling activity of other inner mitochondrial membrane proteins e.g. ANT.

Phenols, benzimidazoles, N-phenylanthranilates, salicylanilides, phenylhydrazones, salicylic acids, acyldithiocarbazates, cumarines, and aromatic amines can induce uncoupling [137]. I anticipate that these will have an anti-cancer activity and merit investigation.

## 2,4-dinitrophenol as an anti-cancer drug

2,4-dinitrophenol (DNP) is a lipophilic anionic (-1) protonophore. It can cross membranes protonated, then lose the proton and return as the anion, then reprotonate and repeat the cycle. It was legally (1933–38, USA), and is now illegally, used as dieting drug. It can be bought over the internet and does cause weight loss [138]. However, the therapeutic margin between a slimming and poisonous dose, which can cause death by overheating [139], is close (3–10 fold) [136]. There were reports of cataracts as a side effect in the 1930s but this relation has been later queried [140]. In the correct doses, the deleterious effects of DNP are few [141]. It has been taken by hundreds of thousands of people, often without proper medical consultation, and typically without ill effect [141]. From 1900–2011 (>100 years), there were 62 deaths in the medical literature attributed to DNP [139]; but some of these were intentional, suicide acts. Although not appropriate as a weight loss pharmaceutical, DNP may be useful as a cancer drug. The risk-reward axis is different for terminal patients – with just days, weeks or months to live – than for healthy adults chasing aesthetic goals. Moreover, a further distinction is that the cancer, with its hyperpolarised Ψ_IM_, will accumulate and sequester the drug from normal cells. Incidentally, low doses of DNP increase lifespan in healthy mice [142].

## DLCs can enhance uncoupling by anionic uncouplers e.g. DNP

The negative DLC and anionic uncoupler will be more targeted to cancer cells because of their more hyperpolarised Ψ_IM_. Thence this enhancement effect, by the DLC upon uncoupling activity [143], will be more targeted to cancer cells. This may be especially pronounced if a DLC with two positive charges is used e.g. dequalinium. If the disparity in Ψ_IM_ is ∼80 mV between normal and cancer cells, for example, dequalinium will localise ∼500 times more to the mitochondria of cancer cells. So, the enhancement effect will be ∼500 times more pronounced in cancer cells. It will permit lower concentrations of uncoupler be used. Relevantly, dequalinium has an anti-cancer activity of its own *in vitro* and *in vivo* [44].

## Drugs that selectively inhibit the reverse, and not the forward, mode of ATP synthase will selectively kill cancer cells

The Ψ_IM_ of cancer cells could be collapsed by a drug that inhibits the reverse, ATP hydrolysing, but not the forward, ATP synthesising, operation of ATP synthase. There are a number of such drugs, based around five different molecular scaffolds (Figure 5), and listed in Table 1 [144]. These drugs may also kill cancer cells by disrupting their pH homeostasis. Unlike normal cells that sit in neutral tissues, cancer cells reside in acidic tumours [120-122] and express ATP synthase at their plasma membrane [145]. They may be using its “reverse” mode to excrete protons. The drugs of Table 1 should be harmless to normal cells as they aren’t using this “reverse” mode of ATP synthase. I urge their trialling on cancer cells and in xenograft mouse models [146]. Especially BMS-199264 [147-149]; it is already being investigated as a protective agent during ischemia [147-148] and can cross plasma and mitochondrial membranes [148].

**Figure 5.**
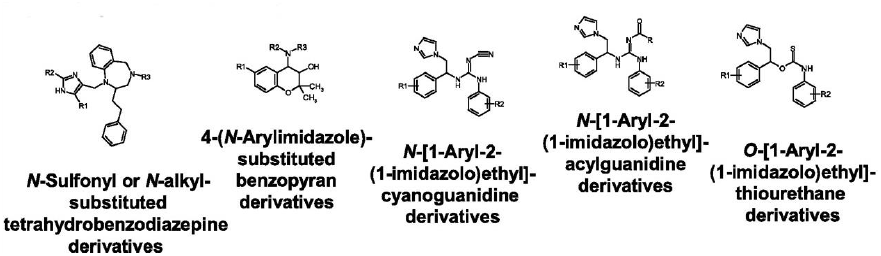
The base structure of drugs that inhibit the reverse, ATP hydrolysing, but not the forward, ATP synthesising, operation of ATP synthase [144]. The R group is variable and some R groups are detailed in Table 1.

**Table 1.**
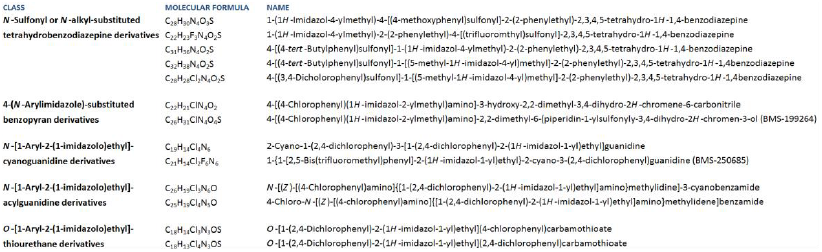
A list of drugs that inhibit ATP hydrolysis, and not synthesis, by ATP synthase [144].

Promoted expression or increased activity of endogenous IF-1 also blocks the “reverse” mode of ATPase and is another direction for anti-cancer drug development. BTB06584 [150] and diazoxide [144] can block this mode through IF-1 and should be tested for anti-cancer activity. Analogues of IF-1, and antibodies of its binding site on ATP synthase, should be explored. Melittin and the 25-residue mitochondrial import pre-sequence of yeast cytochrome oxidase subunit IV (and its synthetic derivatives: Syn-A2, Syn-C, Δ11,12) can act upon ATP synthase like IF-1 [144] and should be tested for anti-cancer activity. Indeed, melittin has anti-cancer activity *in vitro* and *in vivo* [151].

The Ψ_IM_ of cancer cells could also be collapsed by a drug or DNA/RNA that promotes the expression/activity of ANT1, ANT3, UCP2 [78-79]; or a drug/DNA/RNA that decreases the expression/activity of ANT2 [76]. As aforementioned, the (postulated) inverse orientation of ANT2 in the membrane opens many prospects for drug design, wherein the drug is specific to ANT2 (cancer cells) rather than ANT1/ANT3 (normal, adult cells). Drugs may attack the trafficking apparatus that inverts ANT2; if not inverted it would have incorrect activity (prediction). ANT2 expression is under the control of the glycolysis regulated box (GRBOX), which regulates expression of machinery for the aerobic glycolysis operating state [53-54]. This is a possible cancer drug target. These therapies can be utilised in combinations amongst themselves or with other treatments.

These candidate drugs, or DNA/RNA, can be targeted to the negative mitochondrial matrix, and thence disproportionally to cancer cells because their matrix is more negative, by attachment to a DLC. For example, they can be attached to a lipophilic triphenylphosphonium (TPP^+^) cation(s) [152-153], or to its methylated form (TPMP+) or to a VV-arylpyridinium ion [154]. The attachment can be via esterase labile ester bonds that are hydrolysed by esterase enzymes in the mitochondrial matrix, releasing the payload [152]. Or a linkage that is broken by hydrogen peroxide, a ROS present in the matrix [155]. Or it may not be necessary for the linkage to be broken if it doesn’t prevent required interactions. Using dequalinium, or P2C5 or P2C10, as the localising DLC has the advantage that they each have two positive charges and so will be more localised than mono-cations [156-157]. It may be that multiple DLC molecules can be joined together to produce even stronger targeting frames. The more DLC molecules conjoined, the higher the charge of the conjugate and the greater it’s targeting to cancer cells, if it can retain lipid solubility. The lipophilic character may be diminished but this will be equivalent for cancer and normal cells and won’t affect targeting. Multiple TPP^+^ cations have been used relatively unsuccessfully [158] and successfully [158-159] to convey increased targeting.

Atractyloside (ATR), carboxyatractyloside (CAT), epiatractylate (epi-ATR), coffee-ATR, wedeloside, AppCCl_2_p (adenosine-5′-[ß,y-dichloromethylene]triphosphate; a metabolite of clodronate) or ApppI (triphosphoric acid 1-adenosin-5′-yl ester 3-(3-methylbut-3-enyl) ester) are inhibitors of ANT, acting on its “c-side” [81, 160-161]. Antibodies can be produced against elements on the “c-side” and these may act as further ANT inhibitors. These drugs and antibodies aren’t lipid soluble and so may not inhibit ANT2, which I predict is inverted and has its “c-side” in the mitochondrial matrix. However, they may be able to permeate this membrane if they are attached to delocalised lipophillic cations (DLC), conferring them lipophilic character and a net positive charge(s). Perhaps the attachment(s) can be via an ester bond, hydrolysed by esterases in the matrix [152]. Their targeting to the mitochondrial matrix would make them harmless to normal cells, with no relevant vulnerability there, but deadly to cancer cells using an inverted ANT2. If Ψ_IM_ = ∼-140 mV, the *net* positive complex will accumulate ∼225 times more in the matrix than in the cytoplasm and if Ψ_IM_ = ∼-220 mV, as in cancer cells, it will be ∼5000 times more; ∼25 million times more if the complex is *net* double positive (Nernst equation, T = 300 K).

Alternatively these drugs, or others, could be delivered to the mitochondrial matrix by a tiered liposome delivery system. The larger, outer liposome confers passage into the cell of the secondary, smaller, constituent liposome(s) which themselves confer passage past the outer mitochondrial membrane (OMM) for the tertiary, smaller constituent liposome(s) which passage the drug past the inner mitochondrial membrane (IMM) and into the matrix. Only two liposome tiers, rather than three, will be needed if the secondary liposomes can passage the OMM through VDAC channels. The number of required tiers will be cut if the initial, primary liposome is internalized by endocytosis (and so acquires a further bilayer upon cellular entry). Positive molecules can be embedded into the membrane of liposomes to assist in targeting to the negative mitochondrial matrix or a transmembrane potential (negative inside) can be attributed to the liposomes. Targeting molecules can be embedded in liposomes to piggyback upon physiological trafficking processes. Some alternative, tested, and already successful, liposome delivery systems, to mitochondria, are described in [154]. Alternatively, or in addition, these drugs or others could be delivered by attachment to membrane permeant peptides e.g. penetratin or the herpes virus protein VP22 [154, 162].

Antibodies can be produced against elements on the “m-side” of the ANT and these may act as further ANT inhibitors. They likely won’t be lipid soluble and so won’t be able to access and inhibit ANT1 and ANT3, with their “m-side” in the mitochondrial matrix. However, assuming they can passage into the cell (e.g. via liposome entry), they may access the “m-side” of ANT2, which faces the cytoplasm (prediction). So, they would be ANT2 specific inhibitors. They could be attached to a lipophilic anion, which would target them to the mitochondrial intermembrane space. Eosin maleimide (EMA) is membrane impermeable and can only bind the “m-side” of ANT [81]. It could be used in an experiment to test if ANT2 is inverted in mitochondria isolated from cancer cells.

## A drug that inhibits ATP hydrolysis, and not synthesis, by ATP synthase (e.g. BMS-199264) in combination therapy with an uncoupling drug (e.g. DNP)

An uncoupling drug applied with a drug that inhibits the “reverse” mode of ATPase but not its “forward” mode – e.g. BMS-199264 – could bring devastation to cancer cells and leave normal cells unharmed. There would be significant synergy of action between these two drugs. In cancer cells, the uncoupler (e.g. DNP) would act to collapse Ψ_IM_ and BMS-199264 would remove its means to counter this collapse. Crucially, it would lower the amount of uncoupling drug that needs to be given which would aid the safety profile, lowering the risks of the patient overheating. Inhibiting ANT2 would equivalently inhibit the “reverse” mode of ATPase because it would deny it ATP. So, such a drug would also lower the amount of uncoupling drug needed; as would promotors of ANT1 and ANT3 activity.

## An uncoupler in combination therapy with dichloroacetate (DCA)

DCA inhibits pyruvate dehydrogenase kinase (PDK). This decreases PDK inhibition of pyruvate dehydrogenase (PDH) and permits pyruvate to enter the Krebs cycle and OXPHOS to proceed [19]. DCA selectively kills cancer cells *in vitro* and *in vivo* [19], and has caused much excitement [163], but its breakdown products can cause neuropathy [164-166]. DCA acts by constitutively switching on OXPHOS and the ROS produced kills cancer cells [19]. An uncoupler will synergise DCA action by increasing the OXPHOS and ROS production rate. So, it will permit lower DCA concentrations to be used, which will diminish DCA side effects. DCA will reciprocally permit lower uncoupler concentrations to be used (e.g. DNP), which will diminish uncoupler side effects. In a prior paper I proposed exogenous NADH as an anti-cancer drug [34]. I suggest it kills cancer cells as DCA does, by constitutively switching on OXPHOS (by conveying it substrate) [34]. However, unlike DCA, there is likely to be few side-effects as it’s a natural metabolite. It could be used in combination with DCA and/or an uncoupler.

## Conclusion

Cancer cells have a more hyperpolarised Ψ_IM_ than normal cells (∼-220 mV compared to ∼-140 mV). This discrepancy suggests that different processes generate Ψ_IM_ in cancer cells, which may be compromised to selectively kill them. This paper identifies these processes and prospective anticancer drugs, which I hope will be entered into animal and clinical studies. For example, BMS-199264, which blocks the reverse, ATP hydrolysing, but not the forward, ATP synthesising, operation of ATP synthase.

## Funding

The author wrote this paper without any financial support.

